# Trajectories of Response Inhibition Development in Adolescence

**DOI:** 10.64898/2026.04.03.716386

**Authors:** Junda Zhu, Colton R. Smith, Clément M. Garin, Daniel P. Woods, Xin Maizie Zhou, Finnegan J. Calabro, Beatriz Luna, Christos Constantinidis

**Author notes:** These authors contributed equally to this work (Junda Zhu, Colton R. Smith, Clément M. Garin). Corresponding author: Christos Constantinidis < >.

## Abstract

Adolescence is a pivotal developmental period during which cognitive control and the ability to suppress maladaptive impulses undergo profound refinement. Despite extensive behavioral and imaging studies in humans, the neural circuit mechanisms that drive the maturation of inhibitory control remain poorly understood. To address this gap, we performed a longitudinal multimodal study in macaque monkeys spanning the transition from puberty to adulthood, combining behavioral measurements, prefrontal neurophysiology, and structural neuroimaging within the same individuals. Behavioral performance in multiple variants of the antisaccade task improved dramatically across adolescence. In parallel, prefrontal cortical neurons exhibited robust developmental increases in task-related firing, particularly around the generation of goal-directed saccades, indicating maturation of neural representations supporting inhibitory control. Strikingly, trajectories of behavioral improvement and neural activity maturation were tightly predicted by the development of long-range white matter pathways linking frontal cortex with distributed brain networks. These findings identify white matter development as a key systems-level mechanism underlying the emergence of mature inhibitory control and prefrontal neural dynamics and establish a primate framework for understanding developmental vulnerabilities associated with neuropsychiatric disorders that emerge in adolescence and early adulthood.

## INTRODUCTION

The emergence of cognitive control after adolescence is one of the defining features of human development (Larsen and Luna, 2018). Adults acquire an increasing capacity to suppress impulsive behaviors, delay gratification, and flexibly guide actions according to internal goals rather than immediate sensory drives (Mischel et al., 1989; Davidson et al., 2006). Adolescence is characterized by heightened vulnerability to risky decision-making and impaired impulse control, phenomena also linked to increased incidence of psychiatric disorders that manifest during this developmental window, including schizophrenia, ADHD, mood disorders, and substance abuse (Luna et al., 2004; Casey et al., 2008; Van Leijenhorst et al., 2008; Cauffman et al., 2010). Understanding how neural circuits mature to support inhibitory control therefore represents a central challenge in developmental neuroscience with broad clinical relevance.

Behavioral tasks that test response inhibition, such as the antisaccade task (Hallett, 1978), show marked improvement between adolescence and adulthood, providing a robust behavioral assay of cognitive maturation (Kramer et al., 2005; Tervo-Clemmens et al., 2023). Deficits in antisaccade performance are also among the most consistent physiological signatures observed across neurodevelopmental and psychiatric disorders, including schizophrenia (McDowell et al., 2002; Munoz et al., 2003; Smyrnis et al., 2004).

The developmental refinement of inhibitory control parallels extensive structural remodeling of the prefrontal cortex, a brain region critical for executive function (Jernigan et al., 1991; Pfefferbaum et al., 1994; Larsen and Luna, 2018; Friedman and Robbins, 2022). Human imaging studies have documented prolonged maturation of cortical thickness, gray matter volume, and white matter connectivity throughout adolescence (Luna et al., 2001; Gogtay et al., 2004; Ordaz et al., 2013; Bethlehem et al., 2022; Frangou et al., 2022). Functional MRI studies additionally demonstrate developmental changes in prefrontal activation during response inhibition tasks (Padmanabhan et al., 2011; Ordaz et al., 2013). However, fMRI lacks the spatial and temporal resolution necessary to reveal how maturation alters the activity of individual neurons and neural computations that ultimately generate behavior. Consequently, a major unresolved question is how developmental changes in brain structure translate into the maturation of neural activity patterns that support cognitive control.

Non-human primates provide a powerful model for addressing this question because their adolescent development and prefrontal circuitry closely mirror that of humans (Constantinidis and Luna, 2019). Prior work in macaques has shown that response inhibition and prefrontal representations continue to mature after puberty (Zhou et al., 2016a; Zhou et al., 2016c). Anatomical studies suggest a protracted period of monkey prefrontal cortical development that continues during the adolescence period (Lewis, 1997; Xu et al., 2020; Kolk and Rakic, 2022). Cortical thickness, surface area, and white matter myelination also appear to follow similar trajectories to those of humans (Kim et al., 2020; Xia et al., 2023; Alldritt and et al., 2024). Cellular and biochemical changes have also been characterized in the monkey prefrontal cortex from pre-puberty to adulthood, including changes of interneuron morphology and connectivity (Hoftman and Lewis, 2011; Fish et al., 2013). What is still unknown is how structural brain development shapes changes in neural activity that ultimately account for behavioral maturation within individuals over time. Establishing these relationships is essential for identifying the circuit-level mechanisms that drive the emergence of mature cognitive control.

Here, we addressed this challenge through a longitudinal multimodal study tracking adolescent development in macaque monkeys across behavioral, electrophysiological, and neuroimaging domains. We monitored performance in variants of the antisaccade task while simultaneously characterizing developmental changes in prefrontal cortical activity and white matter architecture relative to objective biological markers of maturation. This approach allowed us to directly link evolving cognitive performance with underlying neural and anatomical changes.

## RESULTS

We tracked developmental measures longitudinally in a cohort of four monkeys (Group A, three males, one female) on a quarterly basis, in tandem with neurophysiological recordings and MR imaging, from an age of 3.0 ± 0.1 to 7.1 ± 0.1 years (corresponding to human ages of ∼9-21 years). Another four monkeys matched for training time in the task at different time points were used for control comparisons (group B). To align individual growth trajectories on a biological developmental marker rather than chronological age we relied on the closure of the epiphysial growth plate, a well-established indicator of skeletal maturation in humans (Vandewalle et al., 2014; Kovacs et al., 2022). Thus, we defined a “mid-adolescence” age for each monkey as the time of the tibial epiphyseal closure (see methods). The mean mid-adolescence age across individuals was 4.8 ± 0.3 years, corresponding to a human age of ∼14.5 years, considering that monkeys age roughly 3 times faster than humans (Plant et al., 2005; Herman et al., 2006).

### Performance trajectories

We evaluated working memory performance with variants of the antisaccade task (Fig. 1A-D), which has been used extensively to evaluate response inhibition in animal (Zhou et al., 2016c) and human studies (Tervo-Clemmens et al., 2023). The monkeys were required to observe a visual cue that could appear at one of eight locations and make an eye movement to a location diametric to the visual stimulus. Behavioral performance was collected at time points spaced ∼4 months apart from 3.4 to 6.2 years old. Three variants were used, which differed in their relative timing of the fixation point offset and stimulus onset. Of those, the “overlap” variant (Fig. 1C) was generally the easiest, as it allows subjects to “grasp” their gaze on a visible fixation point at the time of the stimulus appearance, an inappropriate eye movement to which is otherwise more difficult to resist (Zhou et al., 2016c; Zhu et al., 2024).

**Figure 1.**
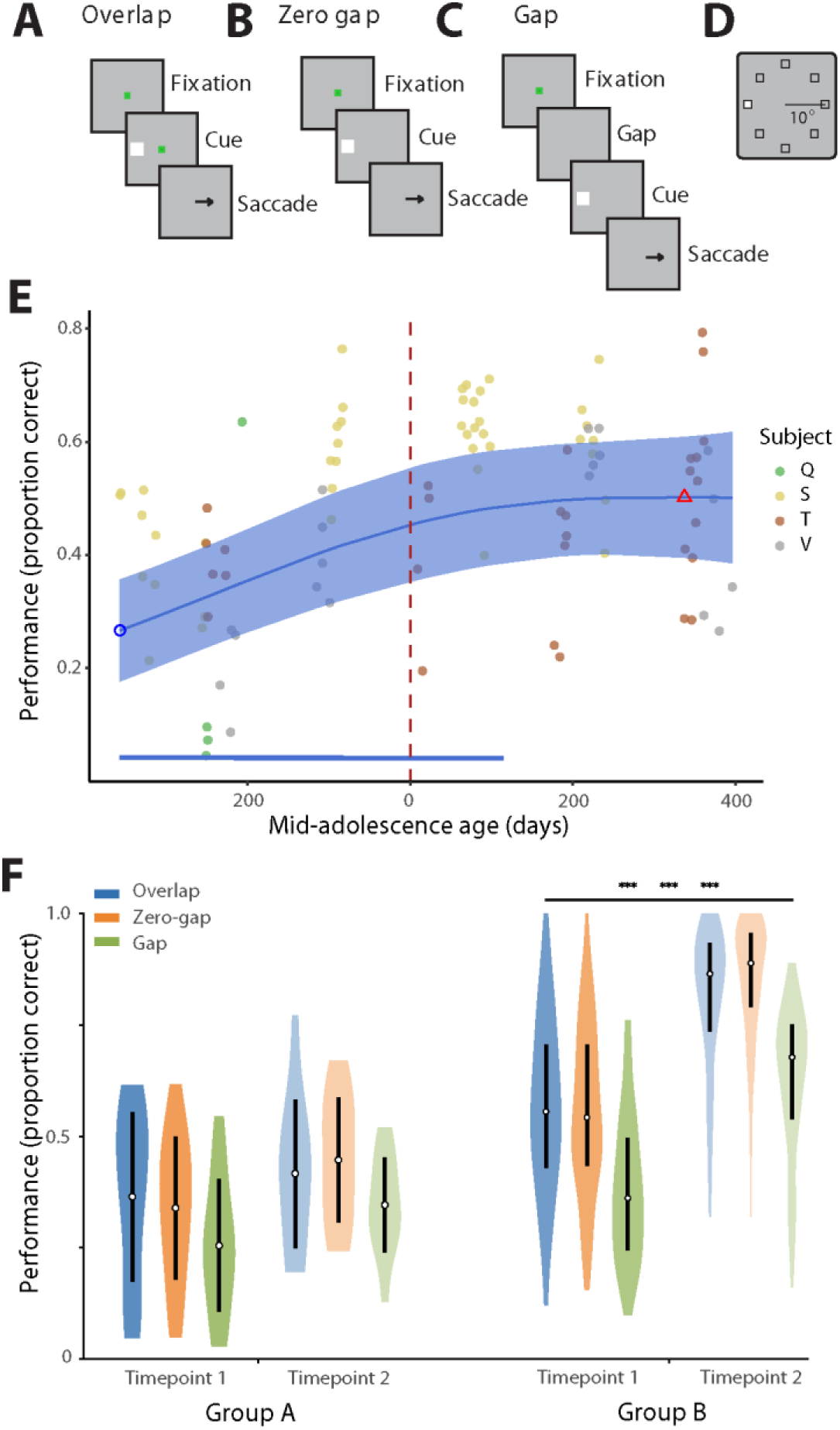
Performance during the antisaccade tasks improves during adolescence. (A-C) Sequence of events in the antisaccade task. The monkey is required to maintain fixation on a fixation point, observe a cue stimulus, and when the fixation point turns off, saccade to a location diametric to the cue. Different task variants were used, in which the fixation point turned off before (A – Overlap variant), simultaneously (B – Zero gap variant), or after the cue appearance (C – Gap variant). (D) Possible locations of the cue stimulus in the screen. (E) Mean performance across all task variants as a function of time, aligned to the mid-adolescence age. Each dot is one session; data from different monkeys are shown in different colors. Blue line shows the GAMM fitted trajectory. Blue circle and red triangle represent minimum and maximum of fitted trajectory. Blue shaded regions denote the 95% confidence intervals (CIs). Dashed vertical line denotes the mid-adolescence age 0. Horizontal bar denotes significant monotonic developmental effect intervals. F. Behavioral performance in the three antisaccade variants in the first two time points for the two cohorts of monkeys (A, tested sequentially through adolescence, and B, for which the second time point of testing was in adulthood).

The levels of behavioral performance in all variants of the task, which we evaluated based on the proportion of correct trials, improved as a function of time with highest gains observed early in adolescence (Fig. 1E). The time course of developmental improvement was similar for the three antisaccade task variants, even though the absolute level of performance differed between them (Supplementary Fig. 1).

The monkeys tested later in development had more cumulative exposure to the task as a consequence of our longitudinal experimental design. To evaluate the effect of exposure to the task, we compared the behavioral performance of this group of animals (Group A) with a group of four animals (Group B, all males) that was introduced to the same task at a similar starting age (median 4.3 years) and were trained under the same protocol (Zhou et al., 2016b). After completing their first time point (young stage), their second time point for behavioral testing began 1.6–2.1years later. We compared performance in the three variants of the antisaccade task between the 1^st^ and 2^nd^ testing time points (Fig. 1F). Group A animals exhibited moderate increases in performance between the first two time points that did not reach statistical significance. In contrast, the performance increase in the second time point for cohort B was highly significant for each of the task variants (Mann-Whitney U test, p<10^-10^ in each case). The result indicates that the difference of age during the two testing time points in two groups accounted for a substantial difference in antisaccade performance between time points.

### Firing rate changes

We recorded single neuron activity from the dorsolateral prefrontal cortex (areas 8a and 46). A total of 2636 neurons were recorded in the four monkeys. To identify the trajectory of activity changes, we first identified neurons with responses to the ODR task, evidenced by a significant increase in firing rate (paired t-test, p<0.05) during any of the task epochs (cue presentation, delay period, saccade), for which correct trials were available in all three antisaccade task variants. A total of 315 neurons (22, 88, 157 and 48 in subjects Q, S, T, and V, respectively) met these criteria. We then divided this sample into three subsamples of equal number of neurons (105 per subsamples) and examined neural activity in the antisaccade task (which was not used as a criterion for neuron selection). The first subsample defined in this manner, spanned -384 to 69 days relative to the mid-adolescent age, the second 70 to 223, and the third 224 to 396.

In agreement with our prior studies (Zhou et al., 2016c; Zhu et al., 2025), we observed that firing rate of prefrontal neurons generally increased across development (Fig. 2A-F). The increase of activity specifically observed in the antisaccade task was evident even after subtracting the baseline fixation rate, and exceeded that observed in the ODR task (Fig. 2G-I). Specifically, we sampled firing rates during the 200ms window after cue presentation and the 200ms window before saccade onset for the early, middle, and late developmental subsamples and subtracted the corresponding baseline firing rate from them. A one-way ANOVA revealed a significant effect of developmental group on firing rate for both the cue period (F_2,312_ = 13.52, p = 2.34e-6) and the saccade period (F_2,312_ = 9.17, p = 1.34e-4). For the cue period, a Tukey post hoc analysis showed that firing rates at the late subsample were significantly greater than those at both the early (p = 1.73e-6) and middle (p = 5.41e-4) subsamples, while firing rates did not differ between the early and middle subsamples (p = 0.42). Similarly, for the saccade period, a Tukey post hoc analysis revealed that firing rates at the late subsample were significantly greater than those at the early (p = 1.51e-4) and middle (p = 3.5e-3) subsamples, while firing rates did not differ across the early and middle subsamples (p = 0.69). Mean activity across all neurons is plotted in Fig. 2; the full distribution of firing rates is shown in Supplementary Fig. 2.

**Figure 2.**
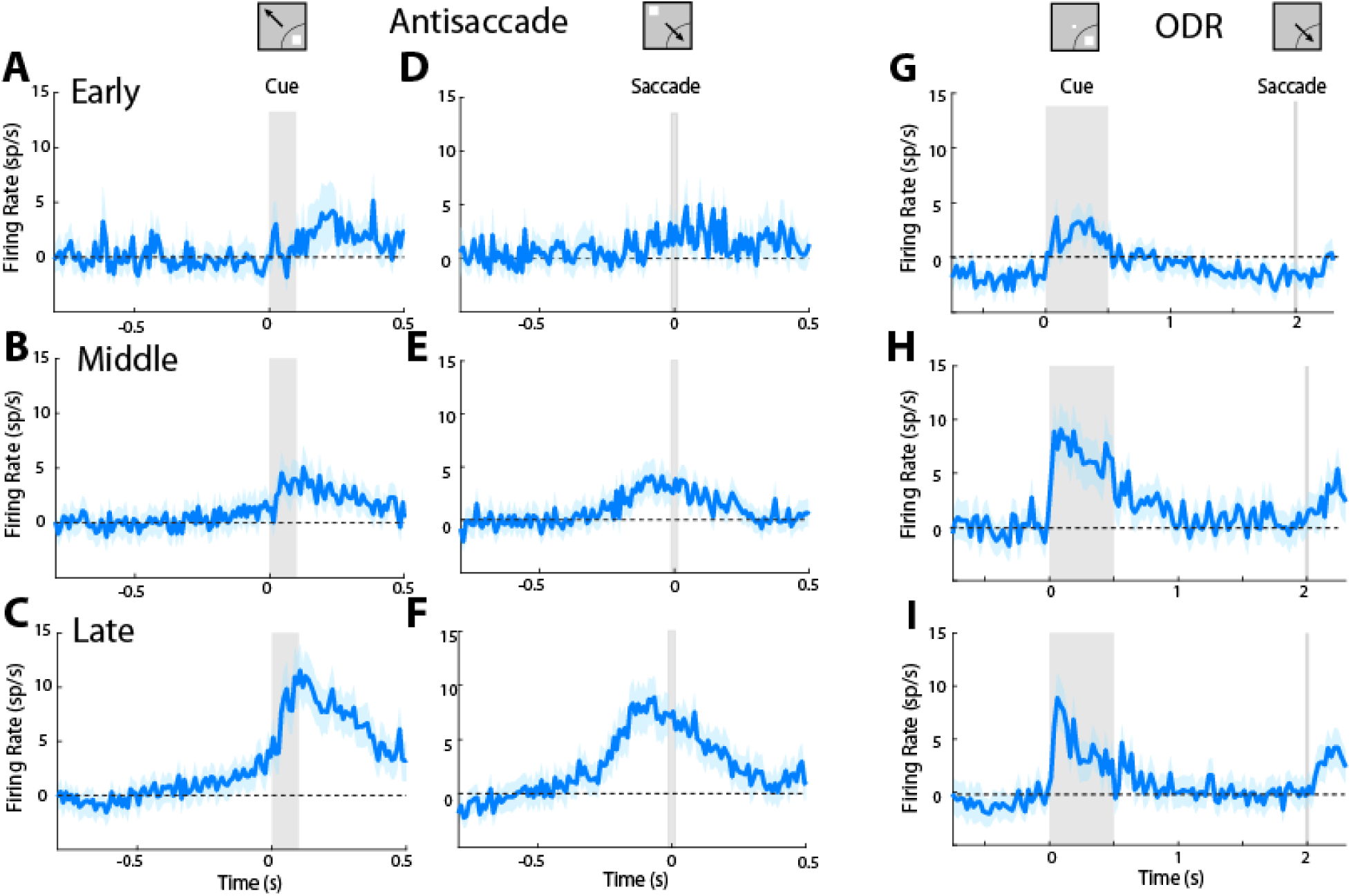
Prefrontal firing rate in the antisaccade tasks increases during adolescence. A-C) Mean firing rate in the antisaccade task, synchronized on cue onset for three successive developmental stages: A, early subsample, n=115 neurons; B, middle, n=115 neurons; C late, n=115 neurons). Bands around the solid line represent the standard error of the mean. D-F) Same as in A-C, for activity synchronized on the onset of the saccade. G-I) Firing rate in the ODR task from the same neurons shown in the antisaccade task. Insets at the top of the figure represent schematically the location of the cue and direction of saccade relative to the preferred location of each neuron used in the population responses, which differed for each neuron.

We have previously described that a critical component of cognitive maturation is vector inversion, the appearance of neural activity representing the location of the saccadic goal, even in neurons that do not otherwise exhibit saccade-driven activity (Zhou et al., 2016c). We thus wished to examine in more detail the trajectory of maturation of vector inversion. We therefore identified visual neurons i.e. those that responded with elevated firing rate during the presentation of the cue stimulus of the ODR task (identified based on a paired t-test, evaluated at the p<0.05 significance level), without exhibiting a significant increase in firing rate during the saccade period of the ODR task. A total of 72 neurons met these criteria (Fig. 3A).

**Figure 3.**
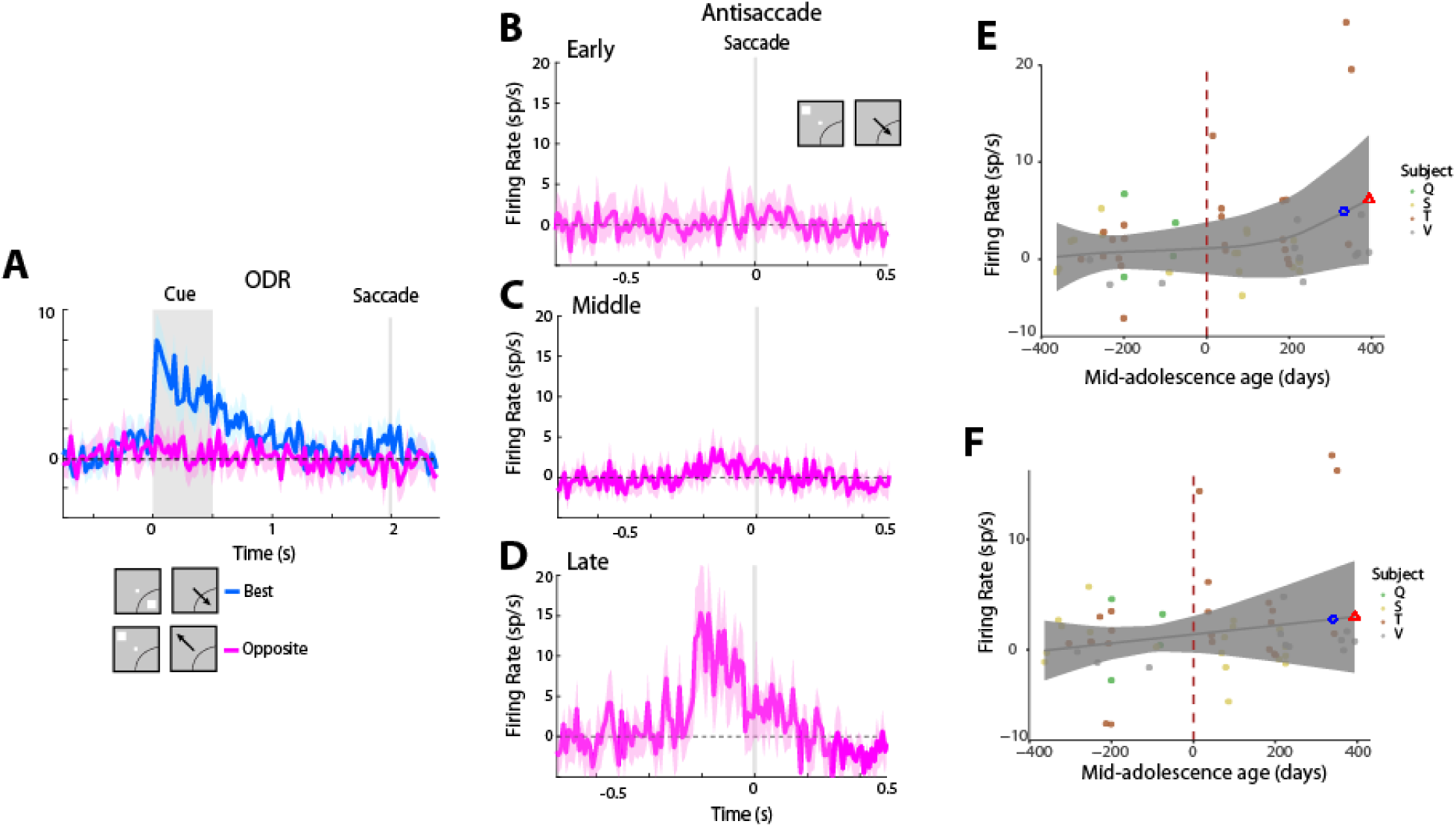
Firing rate of visual neurons. **A)** Firing rate in the ODR task for neurons with only visual activity. Insets at the bottom of the figure represent schematically the location of the cue and direction of saccade relative to the preferred location of each neuron used in the population responses, which differed for each neuron (n=72 neurons). Bands around the solid line represent the standard error of the mean. B-D). Firing rate in the antisaccade task synchronized on saccade delivery, when the cue appeared out of the receptive field for three successive subsamples (early subsample, n=29 neurons; middle, n=33 neurons; late n=10 neurons). Conventions are the same as in Fig. 2. E) GAMM for firing rate aligned to the cue as a function of age relative to mid-adolescence age. F) As in E, for firing rate aligned to the saccade.

We then examined the responses of these visual neurons in the antisaccade task. We specifically focused on the condition involving appearance of a visual stimulus outside of the receptive field, which requires a saccade toward the receptive field, and which would not be expected to elicit a response in neurons with purely visual responses (Fig. 3). The current experiments confirmed that visual neurons were activated in this condition, consistent with the idea that they represent the goal of the saccadic response (Fig. 3B-D). To examine the change in activity for both the cue and saccade intervals more closely, we sampled firing rates in the 200 ms after the cue and 200 ms before the saccade after subtracting the baseline, fixation, firing rate, as outlined above for ODR-responsive neurons. A one-way ANOVA revealed a significant effect of developmental group on firing rate for both the cue period (F_2,69_ = 14.69, p = 4.84e-6) and the saccade period (F_2,69_ = 10.23, p = 1.29e-4). For the cue period, a Tukey post hoc analysis showed that firing rates at the late subsample were significantly greater than those at both the early (p = 6.3e-6) and middle (p = 1.42e-5) subsamples, while firing rates did not differ between the early and middle subsamples (p = 0.91). Similarly, for the saccade period, a Tukey post hoc analysis revealed that firing rates at the late subsample were significantly greater than those at the early (p = 8.9e-5) and middle (p = 7.75e-4) subsamples, while firing rates did not differ across the early and middle subsamples (p = 0.59)

We refined our analysis, by using a Generalized Additive Mixed Models (GAMM) approach, to capture the trajectory of activity and without the potential influence of different baseline activity of different individual monkeys, which may have been sampled unevenly between time points. There was a modest increase in activity both averaged around the time of the cue interval (Fig. 3E) and around the time of the saccade (Fig. 3F). However, the trajectory of activity around the saccade time was more correlated with the trajectory of behavioral improvement (Fig. 1E, r=0.946) than was the cue-aligned activity (r=0.687).

### Relationship with anatomical measures

Having established changes in behavioral performance and neuronal activity during development, we wished to further test what aspects of structural brain changes best explain them. We first examined cortical volume, thickness, and surface, based on generalized additive mixed model (GAMM) fits for the three metrics (Fig. 4). Thickness in the frontal lobe generally increased early in development and reached its peak before mid-adolescence, followed by a decline later in adolescence. In the cohort examined here, we observed a decrease in frontal lobe thickness by −4.7% (Fig. 4). This decrease was particularly pronounced in the lateral PFC −7%. Volume in lateral PFC similarly decreased by −4.5%. Most other frontal cortical regions, including the orbitofrontal cortex and anterior cingulate cortex, did not show significant changes in cortical thickness with maturation (Fig. 4). Similarly, most regions in the other lobes (parietal, temporal, occipital) did not show significant variations (below 3%), except for the medial temporal area, which showed an increase in thickness in our dataset.

**Figure 4.**
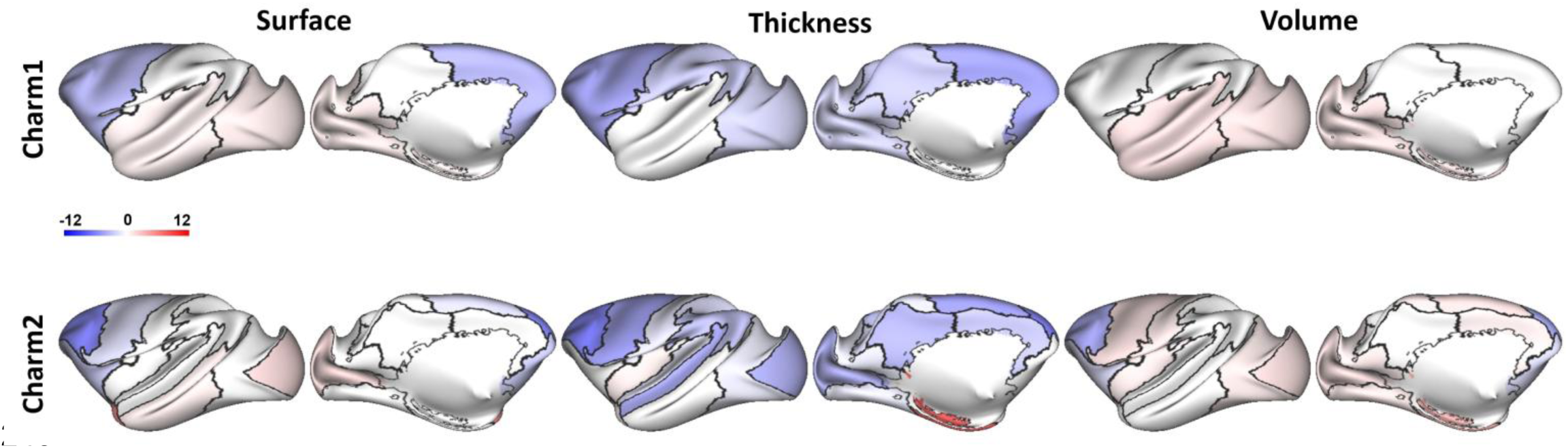
Morphometric developmental changes. Percentage changes in cortical surface area, volume, and thickness during adolescence across brain regions at different levels of segmentation based on the CHARM atlas level 1 (lobar) and level 2 (Jung et al., 2021). Color indicates the percentage change (see color bar) for each region relative to the earliest time points. Changes are projected onto the surface of the NMT template.

We next sought to test how well changes in prefrontal cortical structural metrics predicted the trajectory of behavioral performance in the antisaccade task (Supplementary Fig. 3). Since the performance across all task types was highly correlated with each other, we pooled results from all tasks together. The overall performance trajectory constructed this way was highly representative of the trial-type specific curves (Pearson correlation values of the overall trajectory and individual task trajectories was r > 0.99 in all cases). We observed a modest correlation between the trajectory of behavioral performance and the lateral prefrontal volume (r = -0.798, RMSE= 1.533), surface (r= -0.802, RMSE = 1.451) and thickness (r = - 0.734, RMSE = 1.648).

On the other hand, behavioral trajectories for performance in the antisaccade task matched the maturation of white matter tracts. We examined in detail the white matter tract maturation that most closely paralleled the trajectory of working memory performance improvement (Fig. 5 and Supplementary Fig. 4). We thus performed correlation analyses between Fractional Anisotropy in 55 identified white matter tracts and the behavioral performance of the antisaccade task. The performance in the antisaccade task showed strong trajectory alignment with the FA of most tracts (median |r|=0.9412, median RMSE =0.7083). Examples of high-alignment tracts included the Anterior Cingulum (r=0.9836, RMSE=0.4228) and Medial Longitudinal Fasciculus (MLF; r=0.9673, RMSE=0.4722). In parallel, Radial Diffusivity (RD) showed predominantly inverse alignment with behavior (median r =−0.9387; median RMSE = 0.6771 – Supplementary Fig. 5). These results indicate that white matter maturation showed the strongest trajectory alignment of behavioral improvement during adolescence.

**Figure 5.**
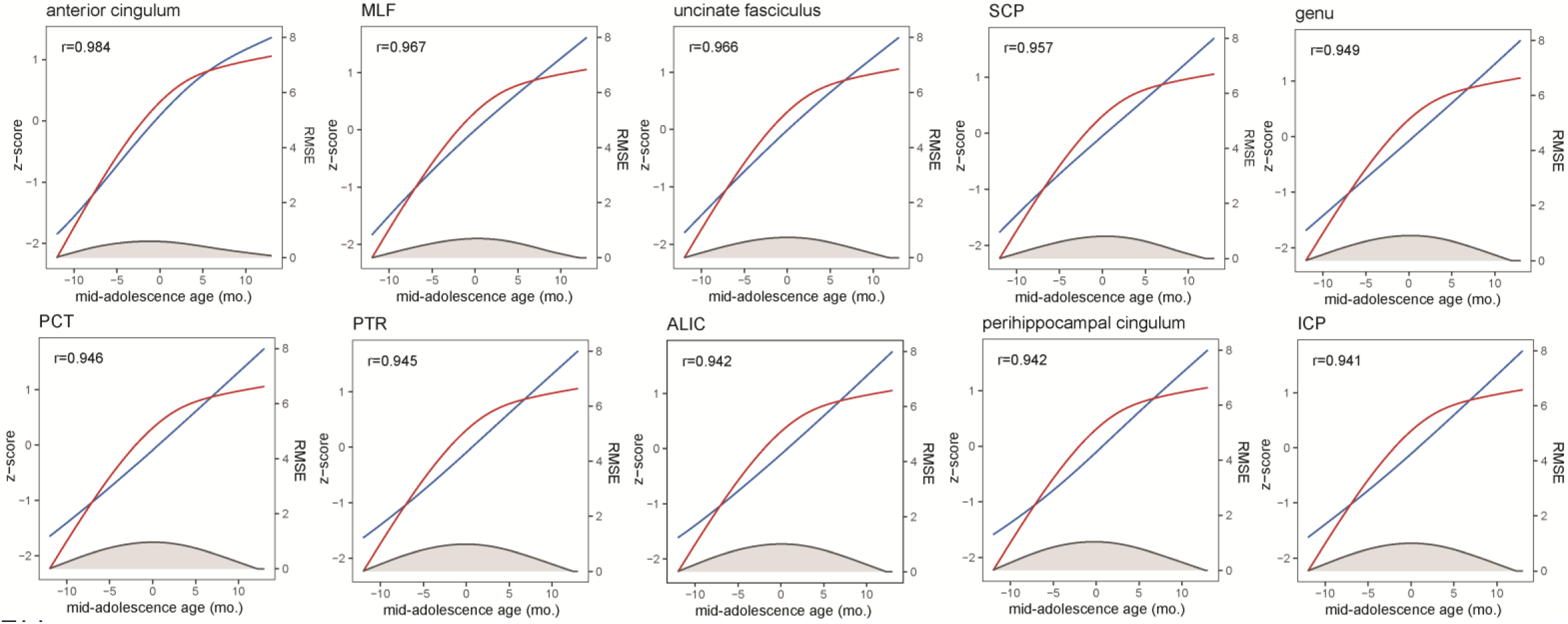
Correlation between performance and FA. Plots of 10 white matter tracks with highest correlation between performance (red curve) and Fractional Anisotropy (FA – blue curve). Shaded area represents root mean squared error.

## DISCUSSION

Our longitudinal study provides direct evidence for coordinated development of response inhibition, prefrontal neural activity, and white matter maturation during primate adolescence. Monkeys, as humans, exhibited substantial variability in development, we therefore relied on objective biomarkers of physical development that allowed us to align growth trajectories of different individuals. We then analyzed multimodal data that allowed us to tie improvements in behavioral to neural activity and structural brain changes. Behavioral performance in several variants of the antisaccade task improved substantially through the adolescent period, accompanied by progressive increases in prefrontal cortical firing rates, particularly around the time of saccade generation. Most importantly, these behavioral and neural trajectories were tightly aligned with the maturation of long-distance white matter tracts linking the frontal lobe with distributed cortical areas. These results identify white matter maturation as a key structural substrate underlying the enhancement of the ability to resist the prepotent influence of an external stimulus during this critical developmental period.

### Maturation of response inhibition

Human cognitive ability improves significantly during adolescence, particularly in domains requiring response inhibition and self-control (Simmonds et al., 2017; Montez et al., 2019). The antisaccade task shows marked developmental improvements from childhood through adolescence in humans (Kramer et al., 2005; Davidson et al., 2006). Deficits in antisaccade performance characterize childhood neurodevelopmental conditions such as ADHD (Munoz et al., 2003) and are prominent in mental illnesses that manifest in late adolescence or early adulthood, such as schizophrenia (McDowell et al., 2002; Smyrnis et al., 2004). Understanding the neural mechanisms underlying the normative development of response inhibition is therefore of considerable basic science as well as clinical importance.

In a recent study, we examined in detail the relationship between reaction time and performance in the antisaccade task (Zhu et al., 2024). Saccades generated within ∼150 ms after the onset of the cue were much more likely to result in errors, as subjects were unable to resist the presence of the cue. Performance rapidly improved as the duration of the saccade became longer. This pattern, which parallels findings in humans (Salinas et al., 2019), reflects a mature capacity to suppress rapid, stimulus-driven responses and execute more deliberate, goal-directed eye movements. Importantly, comparison of performance in adolescent and adult monkeys revealed that adult monkeys improved not only in accuracy but also in reaction time, effectively defeating the typical speed-accuracy tradeoff (Zhu et al., 2024). The observation that saccades generated early after stimulus onset are much more likely to result in errors, as the rapid stimulus transient captures gaze, is consistent with the hypothesis that developmental improvements reflect enhanced inhibitory control over reflexive responses. The progressive increase in ability to resist stimulus-driven capture and generate appropriate delayed responses constitutes a major component of cognitive maturation during adolescence.

### Neural activity substrates of cognitive maturation

In accordance with our prior studies (Zhou et al., 2016b; Zhu et al., 2025) we observed a progressive increase in prefrontal cortical firing rates during development in the antisaccade task that exceeded the changes observed in the control oculomotor delayed response task. The result suggests that the changes are specifically related to the cognitive demands of response inhibition rather than reflecting general developmental changes in motor processing.

A critical component of this neural maturation is the emergence of vector inversion, the representation of the saccadic goal in neurons that do not otherwise exhibit saccade-driven activity (Zhou et al., 2016a; Zhou et al., 2016c). Our current analysis revealed that activity in neurons with purely visual responses (identified based on the pattern of responses in the ODR task following visual stimuli out of these neurons’ receptive fields) progressively increased during development. Furthermore, activity representing the goal of the saccade highly correlated with the trajectory of behavioral performance in the antisaccade task. This result suggests that the improvement in response inhibition is primarily accounted for by a stronger representation of the planned saccade goal, rather than filtering of the cue information, a response towards which needs to be resisted. Fixation period activity tasks also generally increased during the period of development, as we have reported recently (Zhu et al., 2025). While often dismissed as “baseline” or background activity, substantial evidence suggests that neurons active during fixation play a functional role in representing task parameters and suppressing eye movements (Zhou and Constantinidis, 2017; Riley et al., 2018).

How neural activity changes produce behavioral outputs is often opaque, however artificial neural networks have been used successfully to uncover the underlying computations (Masse et al., 2019; Xie et al., 2022). Increases in activity in the antisaccade task have been thus recapitulated in artificial recurrent neural networks, which show increases in unit activity as the network performance on antisaccade tasks improves (Liu et al., 2021). This computational finding suggests that the observed activity changes in neural maturation reflects fundamental principles of optimization in neural systems.

### Structural Brain Development

Recent structural imaging studies have documented trajectories of age-related brain changes across large human cohorts (Bethlehem et al., 2022) and monkey populations (Alldritt and et al., 2024). A hallmark of brain development is the decrease in cortical volume and thickness after childhood (Gogtay et al., 2004). This process is thought to be driven by pruning of infrequently used synapses (Bourgeois et al., 1994), though apparent decreases in thickness may also reflect myelination of gray matter (Natu et al., 2019). Synaptic pruning has been shown to enhance neural computations, leading to enhanced stability and improved efficiency of neural representations (Averbeck, 2022; Liuzzi et al., 2023).

In our cohort, we observed only modest relationship between performance and frontal lobe cortical thickness surface or volume. Our findings instead revealed a striking association between white matter maturation and improvements in both behavioral performance and prefrontal neural activity. Fractional anisotropy in several white matter tracts that interconnect key cortical and subcortical regions involved in eye movement control and executive function showed strong trajectory alignment with antisaccade performance. Radial diffusivity showed predominantly inverse alignment with performance, consistent with the interpretation that increasing white matter integrity underlies cognitive maturation.

These findings align with extensive literature highlighting the protracted maturation of white matter pathways in humans, which continues well into early adulthood (Lebel et al., 2012; Simmonds et al., 2014). Our results suggest that the adolescent period is characterized by a critical phase of white matter maturation that directly enables the integration of distributed cortical networks required for sophisticated cognitive control. The high correlation between white matter measures and both behavioral performance and neural activity trajectories suggests that enhanced white matter integrity facilitates more efficient communication among prefrontal regions and between prefrontal and subcortical structures, thereby improving the neural computations underlying response inhibition.

### Limitations and Future Directions

Some limitations merit acknowledgment in interpreting our findings. Although non-human primates provide the closest animal models to humans, there remain important differences between species that may limit generalization of our conclusions. Specifically, while monkeys undergo developmental trajectories broadly similar to humans, the rate and timing of cognitive development differ between species. Additionally, our neurophysiological recordings were restricted to the prefrontal cortex, motivated by its well-documented role in cognitive control and the specializations of neurons within this region (Zhou et al., 2012; Zhou et al., 2016a; Jaffe and Constantinidis, 2021). However, response inhibition is fundamentally a network-level phenomenon involving distributed circuitry spanning prefrontal, parietal, subcortical, and brainstem regions. The superior colliculus and posterior parietal cortex play a critical role in oculomotor control and inhibition; future work examining how development modulates activity in these distributed networks would provide a more complete mechanistic understanding (Johnston et al., 2007; Qi et al., 2010). Finally, our analysis of neural activity here relied only on single-neuron responses; though changes with increased performance have been described in coordinated neural activation reflected in local field potentials (Wang et al., 2022) and synchronous spiking revealed with methods such as spike-count correlation (Qi and Constantinidis, 2012). Follow up studies can provide a more global picture of factors contributing to response inhibition maturation.

## METHODS

### Subjects

Behavioral, imaging, and neurophysiological recordings were obtained from a total of eight (7 male, 1 female) rhesus monkeys (*Macaca mulatta*). Most data presented here belonged to Cohort A which included four monkeys (3 males, 1 female) and was tracked throughout adolescence. Cohort B included another four monkeys (4 males) and was tested at two time points (early adolescence, adulthood), providing a control for training exposure. All surgical and animal use procedures were reviewed and approved by the Institutional Animal Care and Use Committees of Wake Forest University and Vanderbilt University, in accordance with the U.S. Public Health Service Policy on Humane Care and Use of Laboratory Animals and the National Research Council’s Guide for the Care and Use of Laboratory Animals.

### Developmental markers

We tracked developmental measures of Cohort A monkeys on a quarterly basis before, during, and after neurophysiological recordings. Monkeys of this cohort were first obtained at an age of 2.3 – 2.9 years from a commercial breeding company (Alpha Genesis, Yemassee, South Carolina) where they were mother-reared and lived in species-typical social groups. Once in the laboratory, they were housed in groups of two-three animals, in view of each other, and other conspecifics. To determine each monkey’s developmental progress, we relied primarily on skeletal assessment to best capture physical development (Cheng et al., 2021; Kovacs et al., 2022) and align the growth trajectories of different individuals. Using these measures, we defined a mid-adolescence age for each monkey, defined as the time of each monkey’s distal tibial epiphyseal closure, as observed by veterinary professionals evaluating the X-rays, blind to findings of other aspects of the study.

### Behavioral Tasks

All monkeys were trained to perform the Oculomotor Delayed Response (ODR) Task and antisaccade tasks. The ODR task is a spatial working memory task requiring subjects to remember the location of a cue stimulus flashed on a screen for 0.5 s. The cue was a 1° white square stimulus that could appear at one of eight locations arranged on a circle of 10° eccentricity. After a 1.5 s delay period, the fixation point was extinguished and the monkey was trained to make an eye movement to the remembered location of the cue within 0.6 s. In the antisaccade task, each trial starts with the monkey fixating a central green point on the screen. After 1s fixation, the cue appears, consisting again of a 1° white square stimulus at one of the same eight locations arranged on a circle of 10° eccentricity for 0.1 s. The monkey is required to make a saccade at the location diametric to the cue. The saccade needed to terminate on a 5–6° radius window centered on the stimulus (within 3–4° from the edge of the stimulus), and the monkey was required to hold fixation within this window for 0.1 s. Animals were rewarded with fruit juice for successful completion of a trial. Eye position was monitored with an infrared eye tracking system (ISCAN, RK-716; ISCAN, Burlington, MA).

We used three different variants for the antisaccade task: overlap, zero gap, and gap, differing in the sequence of the cue onset relative to the fixation point offset (Fig. 1). In the overlap condition, the cue appears first, and then and fixation point and cue are simultaneously extinguished. In the zero gap condition, the fixation offset and the cue onset occur at the same time. In the gap condition, the fixation turns off and a 100 ms blank screen is inserted before the cue onset. The monkeys were trained in the antisaccade task with the stimulus appearing at any of the eight stimulus locations also used in the ODR task, however during recordings, four possible cue locations for each condition were used, involving either the cardinal axes or the diagonal axes, so there were 3 x 4 types of trials in each block of trials. The sequence of these 12 trials was randomized in each block. During each recording session, monkeys first performed the ODR task which helped determine the receptive field of neurons recorded online, and then performed the antisaccade task.

The visual stimulus display, monitoring of eye position, and synchronization of stimuli with neurophysiological data were performed using in-house software, and implemented with MATLAB (Meyer and Constantinidis, 2005). We analyzed performance in each variant of the antisaccade task by expressing performance as the overall percentage of trials that resulted in correct responses.

### Surgery and Neurophysiology

The monkeys were initially trained in the tasks mentioned above before their neurophysiological recordings. They were naïve to behavioral training or task execution of any kind prior to the behavioral training. After the animals of cohort A had reached asymptotic performance in the behavioral tasks for the first time, we implanted a 20-mm diameter recording cylinder over the prefrontal cortex of each monkey. Localization of the recording cylinder was based on MR imaging, processed with the BrainSight system (Rogue Research, Montreal, Canada). Recordings in each time point were collected with glass or epoxylite coated Tungsten electrodes with a diameter of 250 μm and an impedance of 4 MΩ at 1 KHz (FHC Bowdoin, ME). Electrode penetrations within the cylinder were placed with a stereotaxic grid system (Crist Instruments, Inc, Hagerstown, MD) to ensure we precisely and evenly sample from the region of interest at each time point, without visiting the exact same grid location, to avoid accumulated damage. Within this grid, neurons were sampled in an unbiased fashion, recording from all neurons encountered in our penetrations, without an effort to select some neurons based on any functional properties. Electrical signals recorded from the brain were amplified, band-pass filtered between 500 Hz and 8 kHz, and stored through a modular data acquisition system at 25 μs resolution (APM system, FHC, Bowdoin, ME).

Recordings were obtained and analyzed from areas 8a and 46 of the dorsolateral prefrontal cortex. Neurons were not pre-screened prior to collection; we recorded from all neurons isolated from our electrodes. Recorded spike waveforms were sorted into separate units using a semi-automated cluster analysis method based on the KlustaKwik algorithm (Harris et al., 2000). Neurons for which at least 4 correct trials in every stimulus condition were available in the ODR task were used in the following analyses.

Neurophysiological recordings were obtained from the monkeys of cohort A at time points spaced approximately 3 months apart from 3.4 to 6.2 years old. Between time points, the animals were returned to their colony and were not tested or trained in any task until their next time point.

### MRI acquisition and preprocessing

Structural MRIs were collected from the monkeys of cohort A every 3 months from 2.8 years (34 months) of age to 5.8 years (69 months) of age. In preparation for the MRI scan, anesthesia was induced using ketamine (5–10mg/kg) and dexmedetomidine (0.015mg/kg), and was maintained using isoflurane. The animals were intubated and artificially ventilated at about 20 breaths per minute. Expired CO_2_ was monitored and maintained between 35 and 45 mmHg. Animals were scanned under isoflurane anesthesia at 1%–1.5%. Heart rate and oxygen saturation levels were monitored using a pulse oximeter. Their body temperature was maintained using warm blankets. The MRI system was a 3 Tesla Siemens MAGNETOM Skyra (Siemens Healthcare, Erlangen, Germany). Anatomical images were acquired using a T1-weighted MPRAGE sequence: TR = 2700 ms, TE = 3.32 ms, inversion time = 880, FOV = 128 × 128 mm, 192 slices of 0.5 mm thickness, resolution = 0.5 mm isotropic. Resting state time series data were also acquired using a multiband EPI sequence: TR = 700 ms, TE = 32.0 ms, flip angle = 52°, repetitions = 700, FOV = 128 × 128 mm, 32 slices, resolution = 2 mm isotropic.

Spatial pre-processing was performed using the EvoDevo NeuroImaging Explorer (EDNiX) pipeline (https://github.com/garincle/EDNiX) as described before (Zhu et al., 2025). This relied on functions from AFNI, ANTs (Avants et al., 2009), FSL (Jenkinson et al., 2012), FreeSurfer (Fischl, 2012) and Connectome Workbench (Marcus et al., 2013) for inhomogeneity correction, spatial and surface registration to a standardized space. A high-resolution NMT template (NIH Macaque Template) as well as the CHARM (Jung et al., 2021) atlas segmentations were registered to the study template. Individual anatomical T1 images (T1^n^) were registered to their T1^last^ and each T1^last^ was registered to the study template. The two movement parameters (T1^n^ to T1^last^ and T1^last^ to study template) were combined to register T1^n^ to the study template. Inversion of these movement parameters was used to register the NMT atlases to the individual T1^n^ images. The volumes (in mm^3^) were calculated using the AFNI function “3dhistog”. Surface and thickness and figures were produced using the Connectome Workbench. Surface, thickness, and volume measurements of cortical and subcortical structures were performed on the hemisphere opposite to the one where recordings were performed.

### DTI preprocessing

Diffusion Tensor Imaging (DTI) data were acquired in pairs with a reversed phase encoding direction in the second scan (e.g., PA vs AP). A diffusion-weighted spin-echo echo-planar imaging sequence was utilized to obtain 82 whole-brain slices of 2mm thickness in 30 directions. Data were processed for analysis using MRtrix3 (Tournier et al., 2019) and the Oxford Centre of fMRI of the Brain Software Library (FSL). The raw DICOM images acquired from the scanner were converted to NIFTI format using dcm2nii, and the corresponding bval and bvec files containing information pertinent to the diffusion gradient were combined across scans. Images were then denoised (“dwidenoise”) and mean b=0 images were calculated. Following this, susceptibility induced and eddy current distortion was corrected using FSL (“TOPUP”) and (“eddycorrect”) respectively. A tensor model was fitted to each voxel (“dwi2tensor”) and fractional anisotropy (FA) maps were calculated (“tensor2metric”). A mask was also created from the T1-weighted image (“bet”), segmenting the brain and non-brain tissue from the whole head.

Each FA image was then registered to the subject’s respective skull-stripped T1-weighted image by affine transformation. These co-registered images were subsequently registered to a diffusion-tensor-based white matter atlas for rhesus macaques (Zakszewski et al., 2014). This allowed for a group analysis of several parameters: 1) the directionality of water diffusion within white matter tissue (Fractional Anisotropy - FA), 2) the mean apparent diffusion coefficient of the diffusion tensor (mean diffusivity – MD), 3) the principal eigenvalue, or diffusion parallel to the principal axis of diffusion (axial diffusivity - AD), and 4) the mean of the two non-principal eigenvalues, or the diffusion perpendicular to the principal axis of diffusion (radial diffusivity - RD). The MD, AD, and RD maps were derived from the diffusion tensor using the same method employed for generating the FA maps. After initial processing, a whole-brain region-of-interest (ROI) analysis was performed, and 55 white matter tracts were selected from the DTI atlas selected, including tracts in the left and right hemisphere, and tracts that were bilateral or crossed the midline. Left and right hemispheres were averaged together to produce a total of 32 distinct tracts analyzed further.

### Generalized Additive Mixed Models

To characterize adolescent development of behavior, activity, and brain structural measures in our longitudinal sample, we used generalized additive mixed models. All GAMM analyses were implemented using the mgcv package for R, each with a smooth function of mid-adolescence age as a covariate, using a thin plate regression spline basis to estimate this smooth function. Random effects in each GAMM included subject-specific intercepts and slopes for mid-adolescence age. For each measurement, we fit a generalized additive mixed model (GAMM) for each outcome variable using mgcv::gamm with a smooth term for maturation: s(mid_adolescence age, k=5, bs=’cs’, fx=FALSE, REML estimation, and subject-level random effects (∼1 + mid_adolescence age by subject). Models were fit using complete cases per outcome and predictor. Predicted trajectories were generated on a fixed maturation grid with 100 grid points. Trajectory similarity was quantified by z-scoring each predicted curve and computing Pearson correlation (r) across the shared grid. Shape mismatch was quantified as RMSE between baseline-shifted absolute trajectories. For those models for which there was a statistically significant fixed effect, the gratia package for R was used to conduct exploratory post-hoc analyses to identify significant periods of developmental change. Specifically, the derivatives of each estimated smooth function of age were approximated using the method of finite differences, and a simultaneous 95% confidence. Because we tested the effect of age on several outcome measures, in separate GAMMs, we applied a false discovery rate (FDR) correction (Benjamini–Hochberg) to control for multiple comparisons. The reported p-values reflect the adjusted significance levels unless otherwise specified.

### Correlation between trajectories

To evaluate similarity between developmental trajectories, we calculated the correlation between the GAMM predictions of different measures. To ensure consistency and comparability, the predictor values (mid-adolescence age) were evenly sampled at 100 intervals between the earliest and latest time points for behavioral data. Each curve was then normalized using z-score normalization.

We calculated two key metrics: the Pearson correlation coefficient (r) and the Root Mean Square Error (RMSE). Before calculating RMSE, each normalized curve was shifted such that the starting point was aligned to zero. This was done by subtracting the starting value of the curve. After shifting, the absolute values of the data points were taken to ensure uniformity in the direction of both curves, facilitating a comparable visualization using RMSE between positively and negatively correlated pair of curves. The RMSE was calculated as:

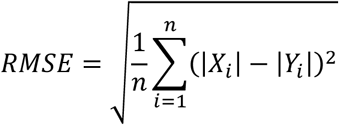

where ∣ 𝑋𝑖 ∣ and ∣ 𝑌𝑖 ∣ are the absolute values of the shifted and normalized data points from the respective curves.

We conducted a permutation test with a maxT approach to assess the statistical significance of the observed correlation coefficients. We randomly permuted 𝑋 and 𝑌 to recalculate the correlation coefficient across 1,000 permutations to simulate the null hypothesis of no correlation. The maxT p-value was calculated to determine the probability of observing a correlation as extreme as the detected one, or more extreme, under the null hypothesis.

The authors report no conflicts of interest.

## ACKNOWLEDGMENTS

We wish to thank Austin Lodish, Leonardo Silenzi, Gracie Hilber, Chrissy Suell, Katelyn Clemencich, and Susan Apt for assistance with experiments; Anna Machado and Adam Anderson for assistance with DTI processing; and Terry Stanford and Emilio Salinas for contributions in the conceptual design of the study. This work was supported by NIH grant R01 MH117996 to C.C.

## AUTHOR CONTRIBUTIONS

C.C. and B.L. conceptualized the project and designed the experiments. J.Z., C.M.G., X.M.Z., performed experiments. C.M.G. and F.J.C contributed to the analysis of the imaging data. J.Z., C R.S. and D.P.W. performed the analysis. C.C. wrote the manuscript with input from all authors.

## DATA AVAILABILITY

Data for the current study will be made available through Zenodo upon acceptance of the paper

## CODE AVAILABILITY

The code used to process the results and generate the figures will be made available at Github upon acceptance of the paper

## SUPPLEMENTARY INFORMATION

**Supplementary Figure 1.**
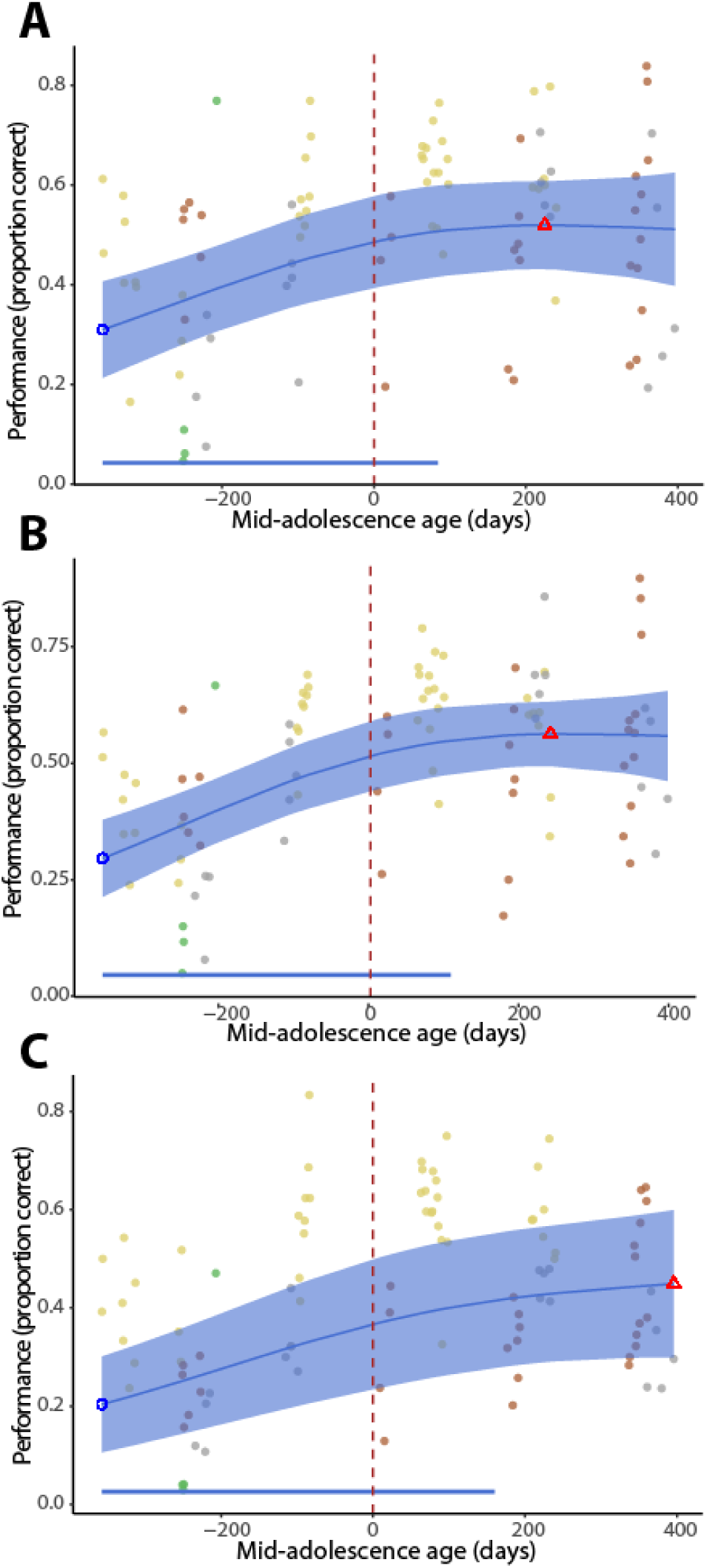
Performance in the three different variants of the antisaccade task.. (A) Overlap variant. (B) Zero gap-variant. (C) Gap-variant. Conventions are the same as in Fig. 1E. Different colors represent the same subjects, as in Fig. 1E.

**Supplementary Figure 2.**
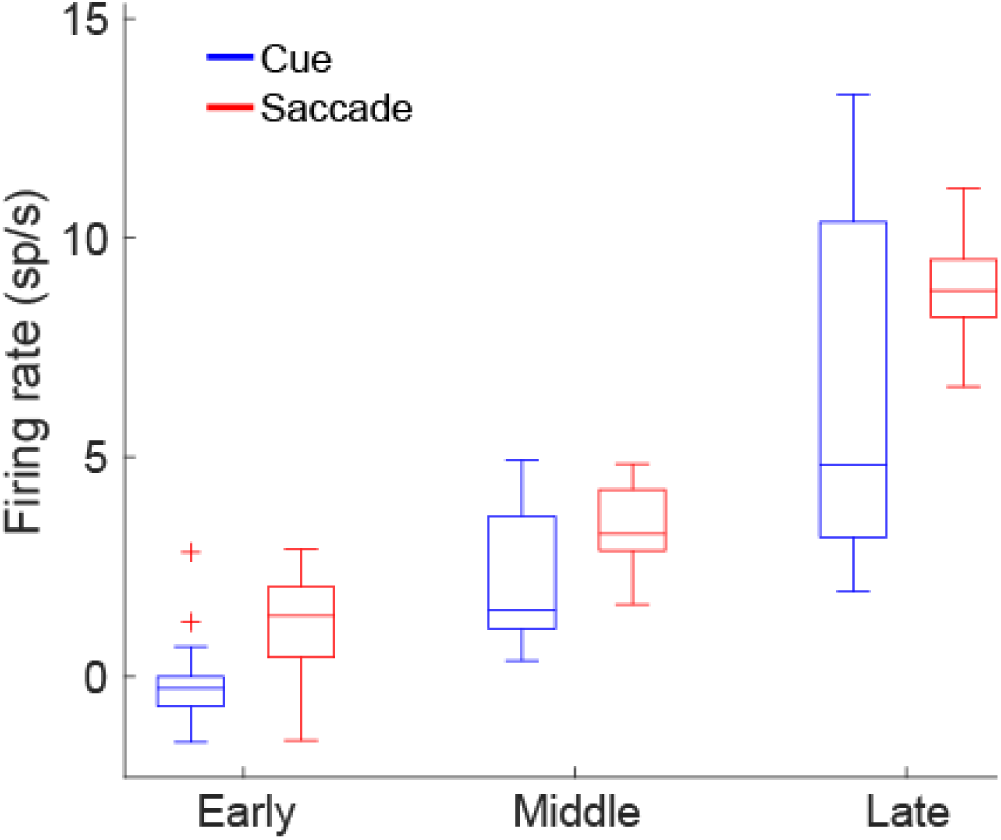
Distribution of firing rates aligned to the cue presentation and saccade, after subtracting the baseline firing rate at each of the early, middle, and late subsamples.

**Supplementary Figure 3.**
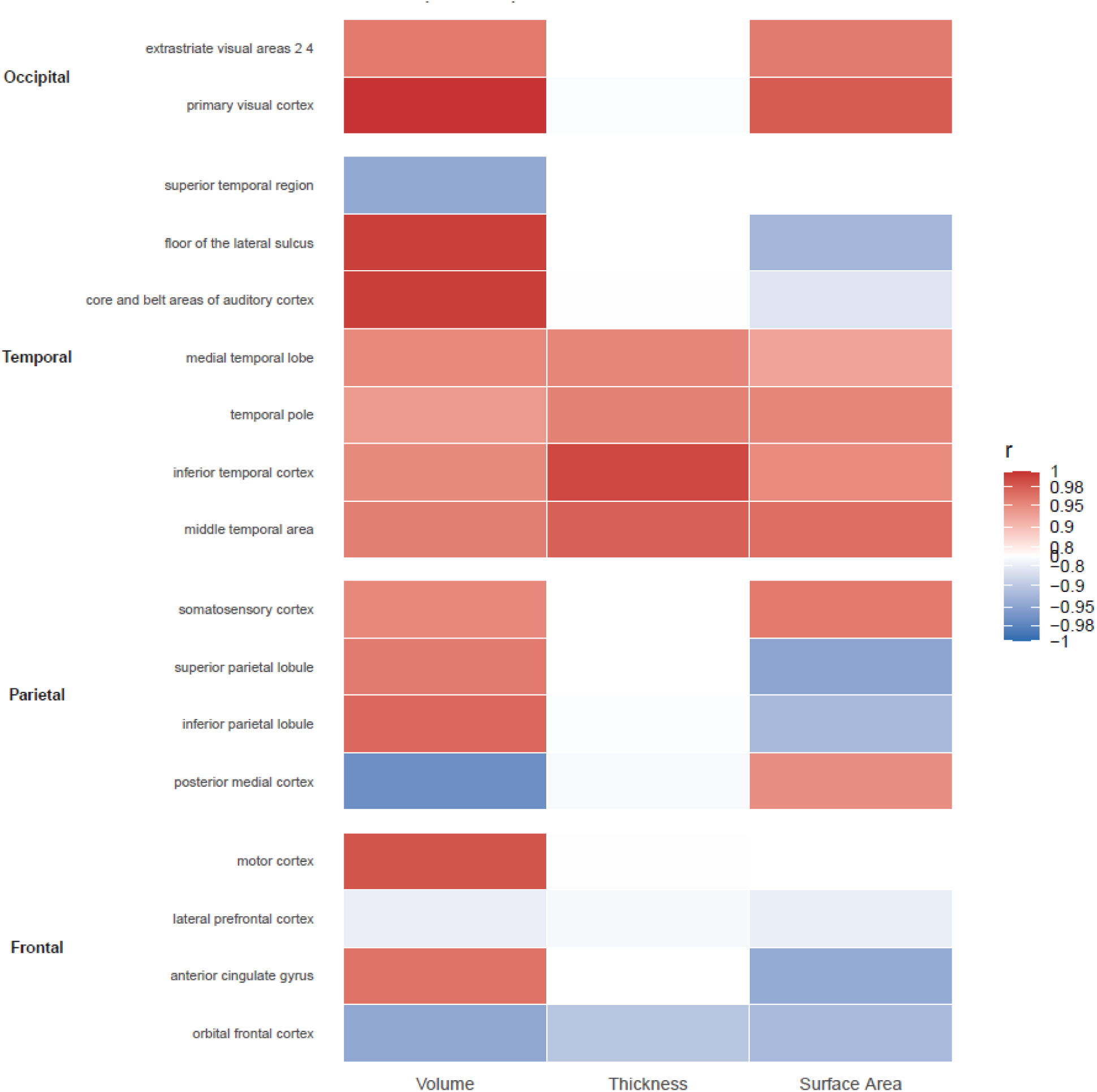
Correlation between behavior, averaged across all task variants, and structural brain measures of volume, thickness, and surface area.

**Supplementary Figure 4.**
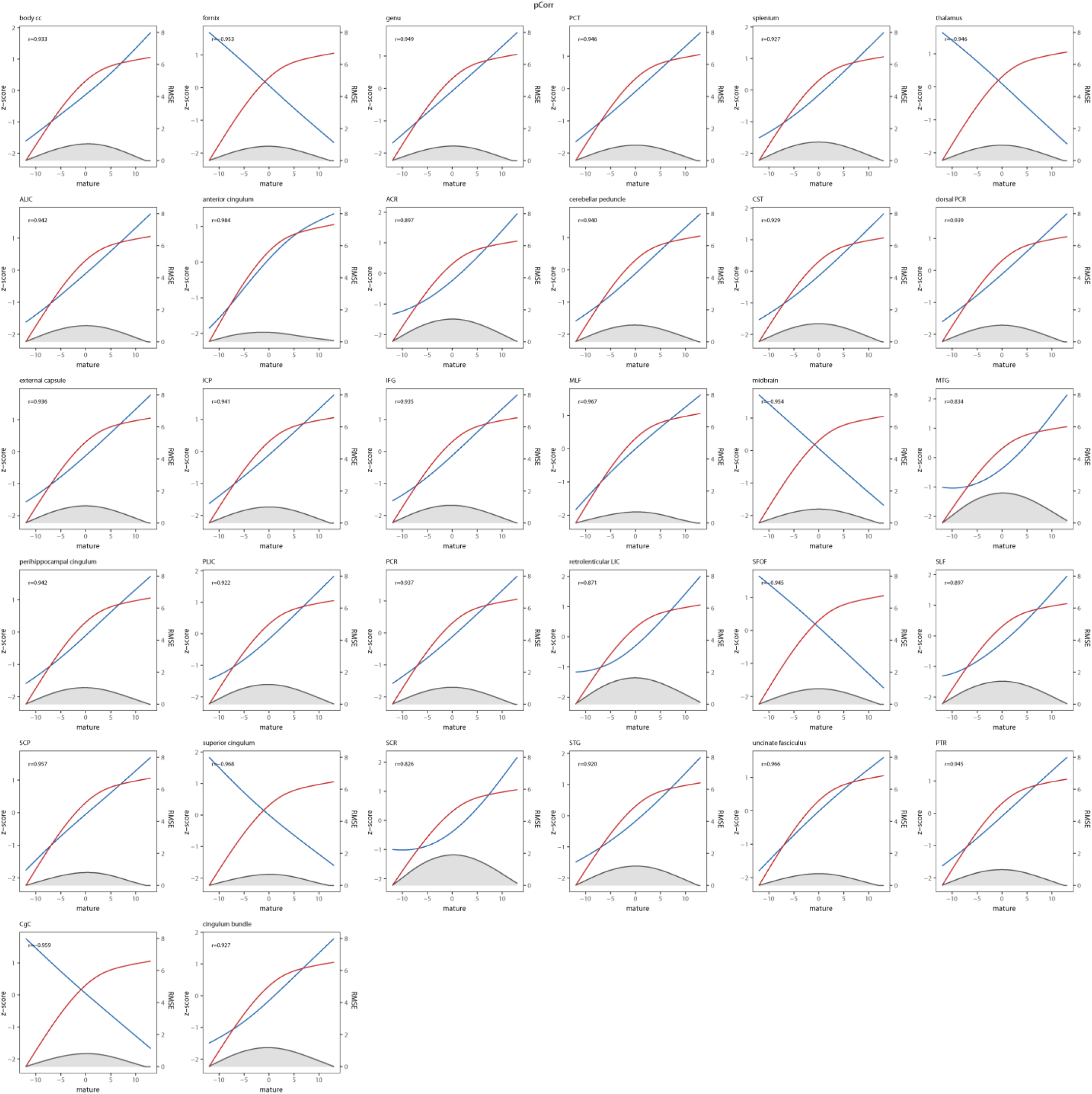
Correlation between performance and FA. Plots of all available white matter tracks showing correlation between performance (red curve) and Fractional Anisotropy (FA – blue curve). Shaded area represents root mean squared error.

**Supplementary Figure 5.**
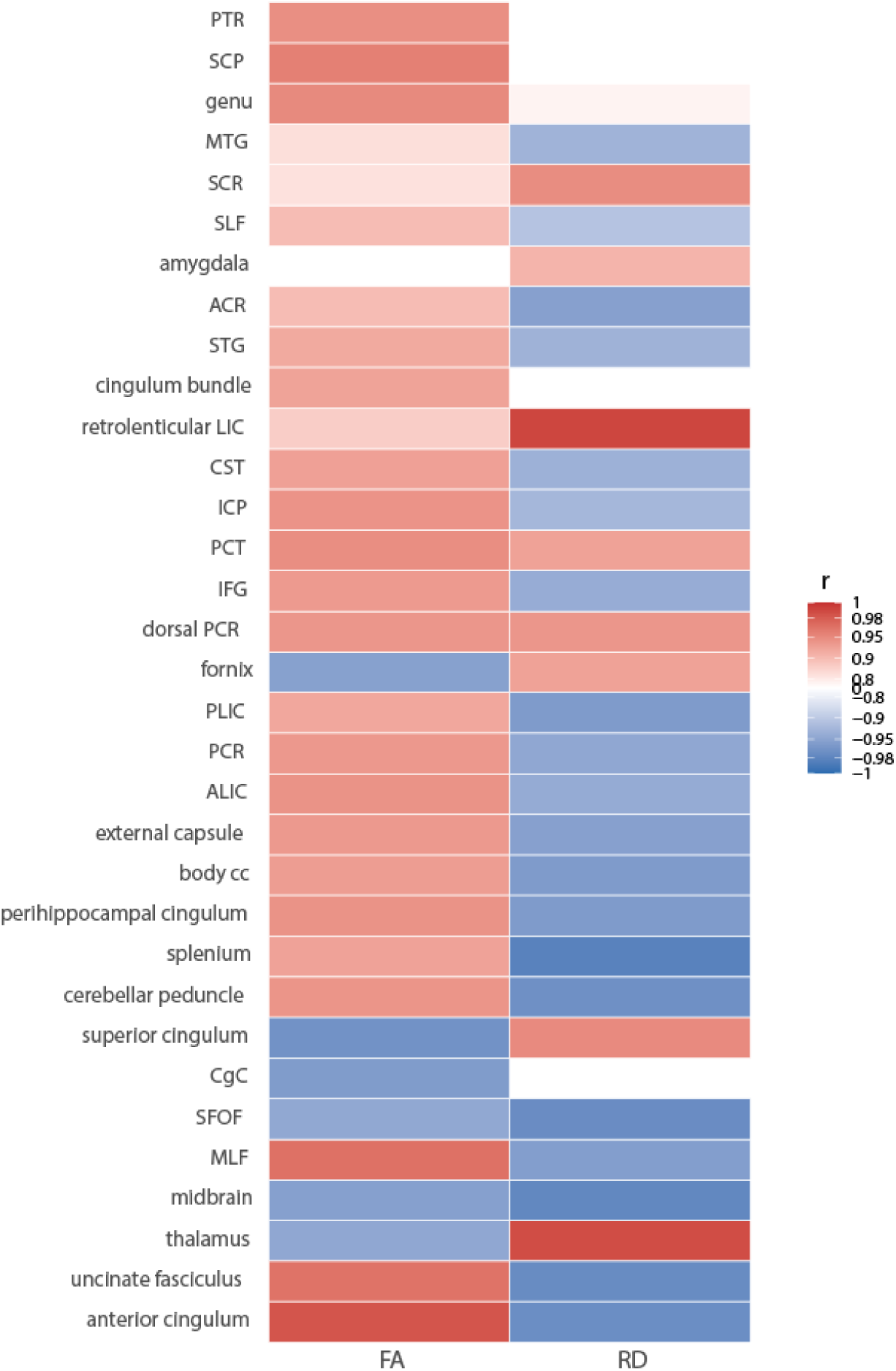
Correlation between behavior, averaged across all task variants, and FA/RD trajectory.

